## Supplemental Table for "A tool for assessing alignment of biomedical data repositories with open, FAIR, citation and trustworthy principles"

### Supplemental material for Murphy et al., 2021

Supplemental table S1

Question identifiers and their abbreviated meaning. PolicyModels allows specifying an id for each question. This id is later used to identify that question, and to localize its text. The ids we use here pertain to the subject of the question. The table below explains each abbreviation.

| **Identifier** | **Subject of question** |
| --- | --- |
| acc | Provide access to the data |
| acc-api | Provide access to the data via API |
| fmt-com | Data format - allow community standards |
| gov-stk | Governance - stakeholders involvement |
| gov-tsp | Governance - transparency |
| land-api | Landing page - machine readability |
| land-ctsp | Landing page - data citation support |
| land-pg | Landing page - pointed by PID |
| lic-cc | License - Creative Commons compliance |
| lic-clr | License - clarity |
| md-FAIR | Metadata - FAIR compliance |
| md-cs | Metadata - using community standards |
| md-daci | Metadata - supporting data citation |
| md-dkn | Metadata - applicability to dkNET community |
| md-level | Metadata - richness level |
| md-lnk | Metadata - linking to publication |
| md-pid | Metadata - includes PID |
| md-prv | Metadata - includes provenance |
| md-psst | Metadata - persistence (even after the data is gone) |
| md-ref | Metadata - qualified references to other (meta)data |
| md-vcb | Metadata - vocabulary usage |
| orcid | ORCID association support |
| oss | Open-Source infrastructure |
| pid-g | PID - using a global PID (e.g. DOI) |
| pid-l | PID - local (e.g. self-assigned) |
| plat | Platform for working with the data |
| reuse | Reuse - licensing |
| ru-doc | Reuse - supporting documentation |
| sch-api | Search and access via API |
| sch-ui | Search and access via UI |
| tr-seal | Core Trust Seal support |

Table S2: Abbreviations used in text

| OFCT | Open, FAIR, Citable and Trustworthy |
| --- | --- |
| FAIR | Findable, accessible, inteoperable and reusable |
| API | Application programmer interface |
| PID | Persistant identifier |
| dk | Digestive, diabetes and kidney diseases |
| dkNET | NIDDK Information Network |
| ORCID | Researcher identifier |
| FORCE11 | Future of Research Communications and eScholarship |
